# DISSEQT - DIStribution based modeling of SEQuence space Time dynamics

**DOI:** 10.1101/327338

**Authors:** R. Henningsson, G. Moratorio, A.V. Bordería, M. Vignuzzi, M. Fontes

## Abstract

Rapidly evolving microbes are a challenge to model because of the volatile, complex and dynamic nature of their populations. We developed the DISSEQT pipeline (DIStribution-based SEQuence space Time dynamics) for analyzing, visualizing and predicting the evolution of heterogeneous biological populations in multidimensional genetic space, suited for population-based modeling of deep sequencing and high-throughput data. DISSEQT is openly available on GitHub (https://github.com/rasmushenningsson/DISSEQT.jl) and Synapse (https://www.synapse.org/#!Synapse:syn11425758), covering the entire workflow from read alignment to visualization of results. DISSEQT is centered around robust dimension and model reduction algorithms for analysis of genotypic data with additional capabilities for including phenotypic features to explore dynamic genotype-phenotype maps. We illustrate its utility and capacity with examples from evolving RNA virus populations, which present on of the highest degrees of population heterogeneity found in nature. Using DISSEQT, we empirically reconstruct the evolutionary trajectories of evolving populations in sequence space and genotype-phenotype fitness landscapes. We show that while sequence space is vastly multidimensional, the relevant genetic space of evolving microbial populations is of intrinsically low dimension. In addition, evolutionary trajectories of these populations can be faithfully monitored to identify the key minority genotypes contributing most to evolution. Finally, we show that empirical fitness landscapes, when reconstructed to include minority variants, can predict phenotype from genotype with high accuracy.

---

Microbial infections, by viruses and bacteria, initially colonize their host as small, quite homogeneous populations, but short generation times and relatively high mutation rates quickly lead to large populations of high genetic diversity. It is well accepted that this diversity facilitates adaptation to the host is through selection of variants from this pool of mutants, in response to environmental change. With the advent of DNA sequencing, viruses and bacteria were the first organisms to be fully sequenced (phage MS2 in 1975^1^; H.influenzae^2^ and M.genitalium^3^ in 1995) and the study of microbial evolution by phylogenetics has benefited from the hundreds to tens of thousands of consensus sequence genomes available for many microorganisms. More recently, High-Throughput Sequencing (HTS) technologies have added new depth to sequence data, capable of quantifying minority variants within the population that differ from the consensus sequence. For example, HTS studies of RNA viruses indicate that both experimental and clinical samples present hundreds to tens of thousands of low-frequency variants, constituting single nucleotide polymorphisms at nearly every nucleotide site along the genome^4;5^. Even before HTS, phenotypic differences between populations with the same consensus sequence have been observed and attributed to suspected differences in variant composition. However, characterization of these mutant ‘swarms’ has generally been limited to mean measures of overall diversity (e.g. Shannon entropy, mean variance, etc.). In a few cases, examples of mixed populations of single nucleotide variations were shown to contribute significantly to virus pathogenesis^6^, fitness and phenotype^4;7^, but focused on only a few variants. Since a mixed population can constitute an evolutionary stable strategy (ESS)^8;9^, the population might aim for an equilibrium where multiple variants coexist.

The rapidly expanding field of single-cell sequencing illustrates how the role of heterogeneity in general can be studied in more and more detail. The data is however complex and noisy, which presents new challenges in the development of algorithms and techniques for analysis, representation and visualization^10–14^. Although phylogenetic tools are well suited for understanding the evolutionary history of lineages and the relationships between lineages/individuals based on whole genome consensus sequence data, they cannot take into account the variant composition hidden by the consensus. Higgins^15^ circumvents these issues by applying multidimensional scaling (MDS) for exploratory analysis, keeping distances between samples more in line with the measured quantities. PhyloMap^16^ superimposes phylogenetic trees on the MDS representation, trying to get the best from both worlds. Relying on consensus sequences only, these models are, however, not well suited for comparison of populations that might be identical at the consensus level but with key differences in the minority variant composition. Other tools are thus needed to adequately represent and visualize a microbial population in sequence space, focusing on where something is, rather than how it got there. Theoretical fitness landscape models, including Wright’s^17^ and the NK landscapes^18^ of Kauffman using two parameters to model the landscape ruggedness paved the way for more recent advances where landscape models are (partially) based on empirical data. One approach is to study the impact of mutations at a few loci only^19;20^, thus artificially enforcing a low dimension of sequence space. To expand the fitness landscape analysis to a higher dimensional setting, Kouyos et al.^21^ utilized predictive models for in-vitro fitness based on the amino acid sequence. For RNA viruses, the mathematical framework provided by the quasispecies theory has been used to describe the population dynamics of these pathogens^22^. Seifert et al.^23^ assumed that viral populations reached mutation-selection equilibrium and applied the quasispecies equation to infer fitness values for the haplotypes in a swarm. However, it is generally accepted that mutation-selection balance is not reached throughout most stages of infection and under most experimental conditions.

Nevertheless, it is tempting to think a proper analysis of an evolving population and its variants composition may foretell whether and where the population will move in genotypic space ^24^, by looking at the dynamics of the population rather than a more static analysis at consensus level. Understanding how the population is developing in sequence space may help predict which directions it can go from there. Viruses, the fastest mutators with small genomes, make ideal model organisms for studying short-term evolutionary processes and will thus be used to showcase the methods developed in this work.

Here, we present DISSEQT (DIStribution based SEQuence space Time dynamics) - a pipeline for analyzing evolution of microbial populations. At the core is a distribution-based model designed to capture the heterogeneity of the populations which makes it possible to describe similarities and differences between populations down to the minority level, and to couple sequence space composition to phenotypic effects. We demonstrate the DISSEQT pipeline with examples from RNA virus and bacterial evolution. First, we show how the DISSEQT sequence space model can uncover biologically relevant features. Second, we followed the evolutionary trajectories of longitudinal samples of experimentally evolved viral populations. Finally, by developing a fitness landscape model based on empirical fitness measurements, we demonstrate how phenotypic effects can be predicted from the population composition. Specifically, we show that the sequence space in which microbial populations evolve are of relatively low dimension, and that biologically relevant signals can be readily captured and used to identify the key variants contributing most to phenotype. We confirm that minority variants contribute significantly to phenotype and must be taken into account for accuracy of genotype-phenotype prediction.

## 1 Results

### 1.1 Overview of the DISSEQT Pipeline

The DISSEQT pipeline (Figure 1, top panel) is designed for reproducibility and openness, from the ground up, using modern software solutions. The source code is openly available in GitHub and all software dependencies are open source. The software can either be installed locally or run directly from Docker images with all required software preinstalled. Running from Docker images simplifies setup and improves reproducibility since differences between local runtime environments are eliminated.

**Figure 1:**
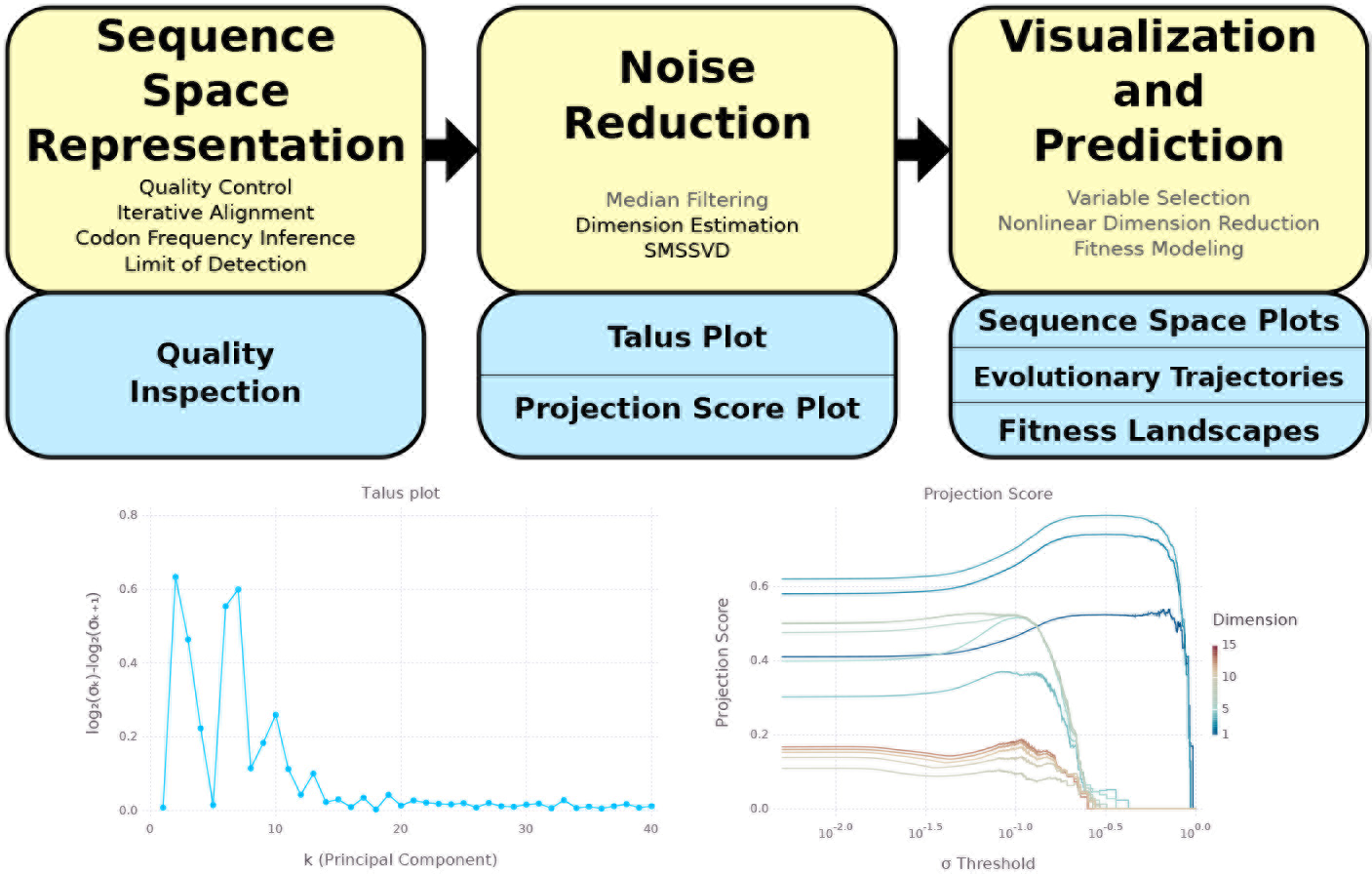
Top: The DISSEQT pipeline. The yellow boxes represent algorithms and data management. The blue boxes represent plots and other output. The analysis history of all results and plots can be traced back all the way to the raw input data. Steps that are only used in some analyses are displayed in gray text. **Sequence Space Representation:** Per sample raw sequencing data is passed through automatic quality control and aligned to a reference genome. Codon frequencies are inferred using quality scores in the aligned data and the limit of detection is estimated for each codon at each site. These are combined to form the sequence space representation. Consensus change reports and read coverage plots aid manual quality inspection. **Noise Reduction:** Median filtering along the time axis is used for time series data. Talus plots are used for dimension estimation and SMSSVD reduces the dimension robustly. **Visualization and Prediction:** Variable selection can be used for finding a small subset of explanatory variables. Nonlinear dimension reduction captures important features for low-dimensional visualization of sequence space. Evolutionary trajectories are described in both sample and variable space. Fitness landscape models are used for visualization and prediction. Bottom left: **Talus plot** for the SynSyn data set. After 13 dimensions, the Talus plot shows small variations around a low mean. Bottom right: **Projection Score Plot** for the SynSyn data set. SMSSVD finds 3 signals of dimensions 3, 5 and 5, with different optima for variance filtering. Each curve displays the projection score of the current signal as a function of the variance filtering threshold, as dimensions are progressively added to the model.

The overview described here is detailed in the Methods Section. The DISSEQT pipeline has three steps, serving different purposes. 1. Establishing a model for sequence space. 2. Reducing noise to make the model robust. 3. Visualization and phenotype prediction.

First, the raw reads for each sample were aligned iteratively until the consensus sequence converged and both automatic and manual quality controls were performed. Maximum likelihood estimation was used to infer the codon frequencies for each position, using all reads overlapping that position, based on a multinomial model with noisy observations. An initial sequence space representation was then constructed using the codon frequencies and a limit of detection estimated for each possible variant at each site. In this article, we focus on coding regions, which makes codons the natural basis for sequence space modeling, since they are closely connected to biological function and this choice does not impose any assumptions about the relative importance of synonymous versus nonsynonymous changes. All methods presented here are also applicable to non-coding regions, by basing the sequence space model on nucleotides rather than codons.

Second, a dimension estimate of the data was obtained by generating a Talus plot (Figure 1, bottom left panel, and Supplementary Methods), after which noise reduction was performed by SubMatrix Selection Singular Value Decomposition (SMSSVD)^25^. SMSSVD is ideal for situations where complex data containing a very large number of variables have signals spread out over different (possibly overlapping) subsets of variables, with the goal of recovering all signals that can be detected, rather than only the strongest one.

Finally, the resulting sequence space representation was used for visualization and phenotype prediction. The evolutionary trajectories of viral populations were followed through time, using sparse methods to find low-frequency variants arising and driving the movement in sequence space. Empirical fitness values were used to create fitness landscapes for prediction, using the representation from step 2, and for visualization, after an additional nonlinear dimension reduction step vital for getting a useful representation in 2d. The sequence space model created by the DISSEQT pipeline is also intended to be used as input to other software packages, e.g. for clustering and regression.

### 1.2 Generation of synthetic synonymous viral lineages with altered localization in sequence space and different minority variant compositions

Our goal was to develop and evaluate a pipeline that can capture the discrete signals within the swarms of variants in clinical or experimental samples - essentially, to monitor and analyze evolving populations before significant changes to consensus sequences occur. To do so, we generated a collection of samples that would be representative of such populations, bearing differences in minority variants. We used four genetically trackable virus populations that derived from the same infectious clone wild type Coxsackie virus B3. Within the capsid-coding region of wild type virus, 117 Serine and Leucine codons are represented by all six codons for each amino acid. We generated three additional synthetic synonymous (SynSyn) virus lineages (Figure 2), some of which were previously published^26^, in which these 117 codons were changed to belong exclusively to only one of three codon categories. These lineages were designed to retain the initial functional neutrality (that is, same protein sequence), while occupying different starting points and potentially different trajectories in sequence space. Indeed, the differences in fitness values and phenotypes are small in comparison to the differences we observe within the lineages in the experiments described below (see Supplementary Figure 1). However, the lineages should behave differently as mutations accumulate at these codons, by accessing different mutational neighborhoods with differing impacts of virus fitness.

**Figure 2:**
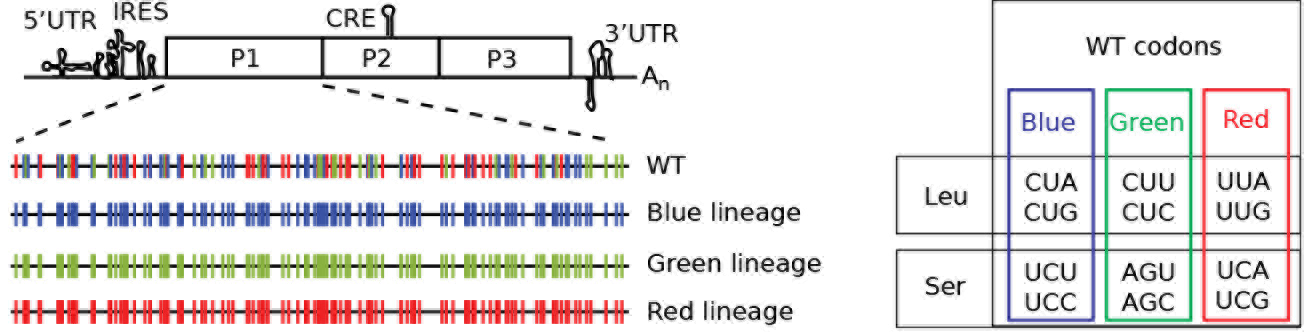
Right: Clusters of Leu/Ser codons according to different viral lineages. Color coding corresponds to synonymous codons used to genetic engineered each viral lineage ‘Blue Lineage’ (blue), ‘Green Lineage’ (green) or ‘Red Lineage’ (red). Left: Schematic of the Coxsackie virus genome indicating RNA structures required for replication (5’UTR, IRES, CRE and 3’UTR) and the single open reading frame encoding capsid structural proteins (P1 region) and non-structural proteins (P2, P3 regions). The P1 region, in expanded view, shows 117 Ser/Leu codons for the wildtype (WT), Blue, Green and Red viral lineages.

Next, to introduce changes in minority variant composition without significantly altering their consensus sequences, we evolved these virus populations in different conditions. Wild type and SynSyn viruses were serially passaged five times in triplicate in normal conditions, as well as in five different mutagenic conditions that are known to increase this virus’s mutation rate ^27^ three base analogues (ribavirin, 5-fluorouracil and 5-azacytidine), amiloride, and Mn^2+^. Low to moderate concentrations were used to accelerate evolution, while higher concentrations were employed to exacerbate fitness effects. We thus obtained 411 mutant swarms (301 passing strict quality controls) from these varied growth conditions, which were deep sequenced to obtain their entire variant compositions. Importantly, passaged samples in each lineage did not have significant consensus changes (in total across the samples, 144 substitutions at 4 different receptor binding sites and 35 substitutions at 12 other sites).

### 1.3 DISSEQT reveals that the sequence space occupied by evolving microbial populations is of intrinsically low dimensionality

Theoretical sequence space is incredibly large, even for a small genome of length *n* = 10000, the number of possible sequences are 4^*n*^ ≈ 4 · 10^6020^, so large that the number of atoms in the universe is miniscule in comparison. The number of sequences reachable within just *K* = 10 mutations, 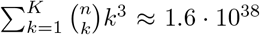 is still vast, and it is unknown much of sequence space is occupied by an evolving microbial population. We generated Talus and Projection Score^28^ plots from the sequence data, which provide a visualization of how the contents of a data set spread out across different dimensions. These plots provide a qualitative estimate of the number of dimensions needed to capture all biologically relevant signals that stand out above the background noise. As shown in Figure 1, bottom left panel, the Talus plot settles after 13 dimensions, with small variations around a low mean, giving a dimension estimate of 13. In the Projection Score plot (Figure 1, bottom right panel), SMSSVD has detected three signals, of dimensions 3, 5 and 5, where the variance filtering threshold for automatic noise reduction has been optimized for each signal.

Next, we examined which biological signals were captured in each dimension and whether analysis of minority variants could better monitor the evolving populations compared to consensus sequence analysis. To do this, sequence space representations of the mutant swarms were generated after noise reduction, where the final SMSSVD step decomposed the samples by principal components. Since almost no consensus changes occurred during the experiment, the principal components found patterns essentially related to differences in minority variants between mutant swarms. As shown in Figure 3, the strongest signal, described by the first three principal components, clearly separates the samples in sequence space according to lineage (see rows 1-3, above the diagonal, in Figure 3). Importantly, further analysis of lower dimensions identified all biological treatments that were imposed on the viral populations. A complete separation in sequence space was observed for mutagenic treatment by 5-fluorouracil, ribavirin, and 5-azacytidine (see rows 4, 5, and 7 below the diagonal, in Figure 3), known to introduce specific nucleotide substitution biases. Even for treatment with Mn^2+^ and amiloride, which increase natural mutation rates without introducing nucleotide bias, a biological signal could be identified in most of the mutant swarms separating from other samples in rows 9 and 11 (Figure 3). Furthermore, these signals are detected despite the background noise and error introduced by the sample preparation and sequencing technology, which lies in even lower components. Finally, if the same analysis is performed using only each sample’s consensus sequence, barely any biologically relevant signals are detected and no patterns related to the mutagens are found (Supplementary Figure 2). These results reveal an important feature of evolving microbial swarms: despite sequence space being of theoretically ultra-high dimension, we showed here that evolving microbial populations such as RNA viruses, which present the highest mutation frequencies, are of intrinsically low dimensionality. Indeed, all five of the biological pressures placed on these viral populations could be captured within the first 13 components.

**Figure 3:**
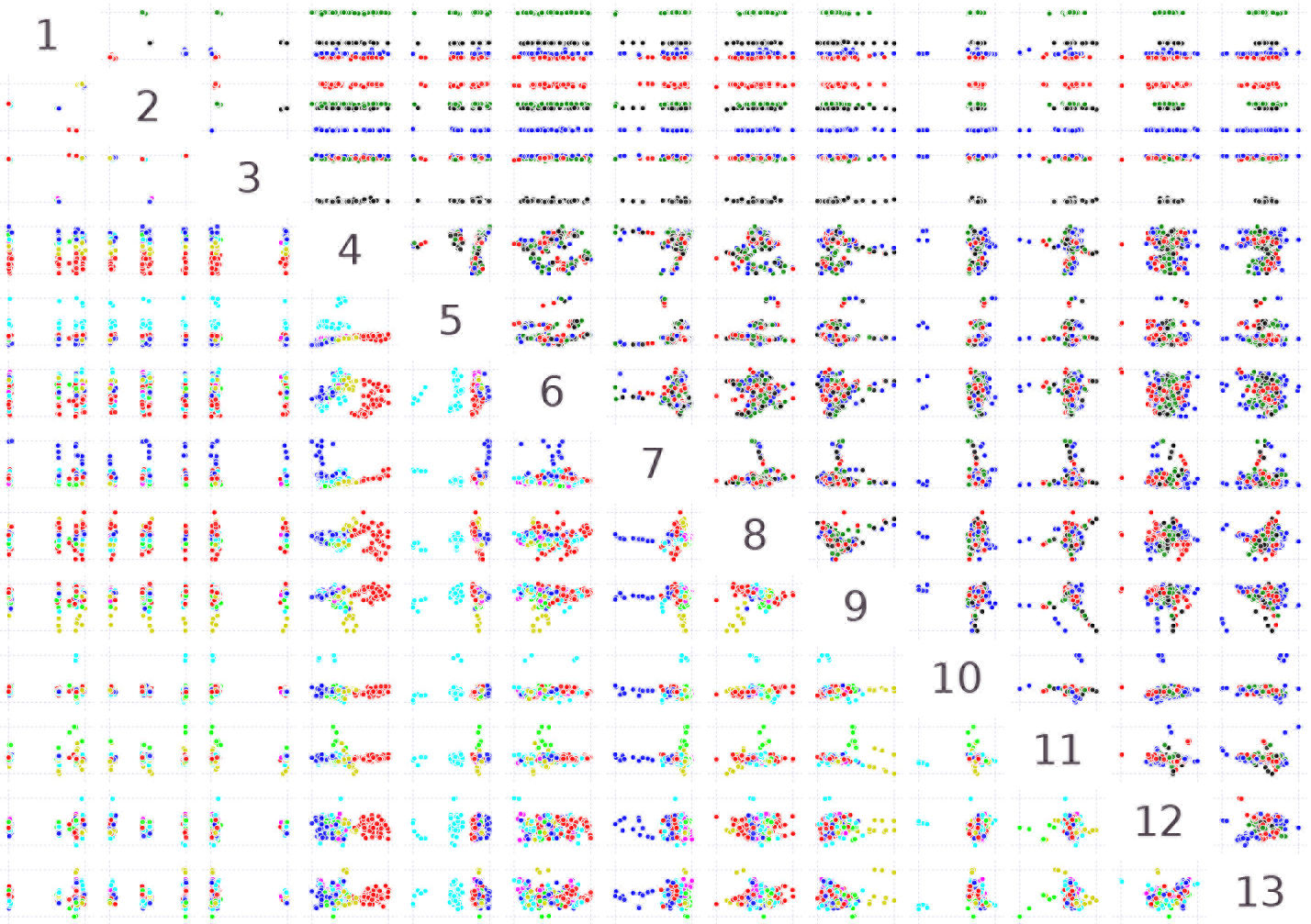
Pairwise scatter plots showing the first 13 principal components in the analysis of the SynSyn data set plotted against each other. Plots above and below the diagonal are mirror images of each other. Each dot represents one viral population. Above the diagonal, samples are colored by lineage 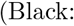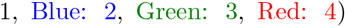 and below the diagonal, samples are colored by mutagen 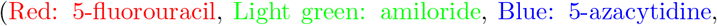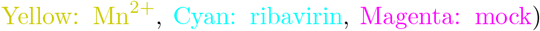. All axes are rescaled to fill the plot area.

### 1.4 DISSEQT can monitor evolutionary trajectories and identify the minority variants involved in adaptation

Recently, we studied the adaptation of Coxsackie virus to a new cell line. Long term passages of experimentally evolved populations (120 generations per virus) were analyzed by deep sequencing. Lacking suitable computational tools, the original study focused on identifying variants in the structural protein-coding region of a wild type lineage that showed signs of positive selection in the final passages of adaptation (mutations appearing at > 2% in more than one replicate, and only in the structural proteins known to be involved in adaptation to cell culture) ^4^. In that study, we identified one consensus sequence change that occurred in all lineages during the first 10 passages, followed by a cluster of minority variants that reached above 5% in the last passage in each series. The data set however, contained whole-genome sequencing for three lineages of this virus: wild type, a higher replication fidelity lineage and a lower fidelity lineage. Using DISSEQT, we could obtain a more complete picture by monitoring the evolutionary trajectories of three biological replicates per lineage (Figure 4), without biasing towards non-synonymous mutations in the structural protein region. The top panel gives an overview based on nonlinear dimension reduction, showing how the evolutionary trajectories of the replicates relate to each other. For each pair of replicates, the time of bifurcation was computed and this was extended to sample clusters using average linking hierarchical clustering. Before the time of bifurcation, the replicates are close in sequence space and follow the same evolutionary trajectory. The splits in the panel show when the bifurcations occur. All replicates shared the same starting point. Around passage 4, the low fidelity replicates (yellow-orange) split from the others and shortly thereafter (around passage 5) the wildtype replicates (magenta-purple) split from the higher fidelity replicates (green-cyan). These observations reflect what was expected, but could not be detected using classical approaches that monitored only a few positively selected alleles: that low-fidelity, mutator strains generated more minority variants more rapidly compared to wildtype, and to high fidelity strains. The replicates within each lineage then followed similar trajectories until further bifurcating between passages 7-19. As with the previous examples where lineages clustered together, these results also support the notion that although sequence space is theoretically huge, similar lineages will tend to travel along the same evolutionary trajectories during the initial periods of evolution.

**Figure 4:**
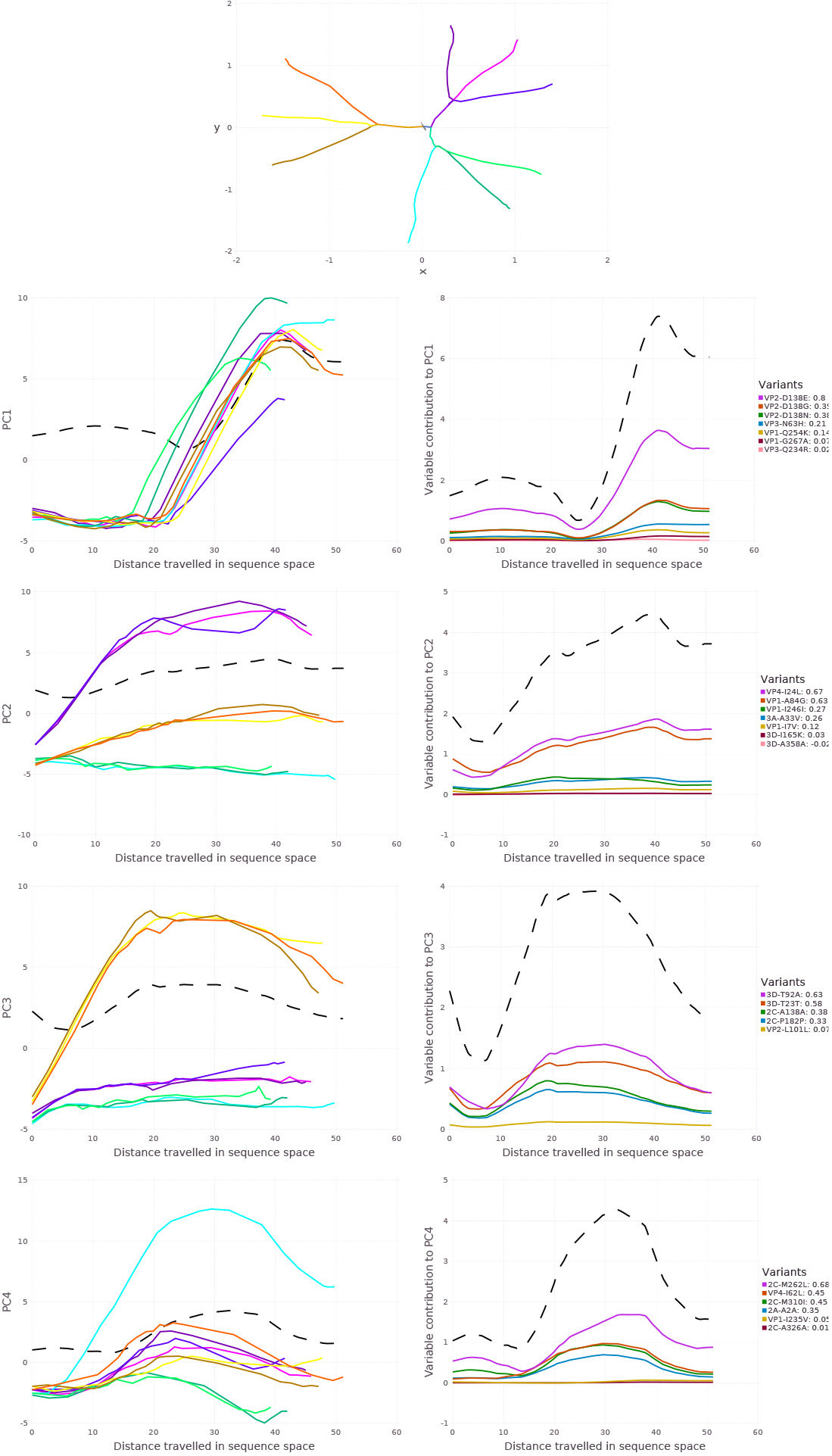
Top: Overview of the evolutionary trajectories of the 9 replicates in the adaptability data set^4^, shown after nonlinear dimension reduction. WT replicates are shown in magenta-purple colors, replicates from the high fidelity lineage in green-cyan colors and replicates from the low fidelity lineage in yellow-orange colors. The starting point in sequence space is very close for all replicates. The splits indicate when the evolutionary trajectories bifurcate, i.e. when the replicates start to deviate from each other. Left column: Principal components for replicates as a function of arc length. Right column: Variable contributions as a function of arc length. Both columns: The dotted black line shows the total contribution to *σ*_*k*_ at *s*.

While the above analysis gave information on rate and direction of evolution, it did not identify what minority variant component of each population was responsible for adaptation and the observed evolutionary signals. Thus, we broke the analysis down by component over time after variable selection, where principal components determined by SMSSVD followed by SPC maps the trajectories of each replicate (left panels), and identifies which variants contributed most to the signal in each component (right panels). The strongest signal in the first principal component captured time dynamics shared between all replicates regardless of lineage, which consisted of the amino acid residues in the structural proteins responsible for adaptation to receptor usage^4^. The remaining components, however, identified several other mutations at sites that were missed by using the classic cut-off of 1-2% minority variant frequency and that could explain differences subtler phenotypic differences between lineages and between replicates. For wildtype, for example, two additional amino acid changes in the VP1 and VP4 structural proteins contributed most to these lineages’ departure from others (principal component 2). Finally, the lower components (4 and onward) revealed variants that explain each replicate’s divergence from others, including many variants in non-structural proteins such as the 2C (helicase) and 3C (protease) (Supplementary Figure 3). Together, the results show that while low-frequency variants were identified at nearly every nucleotide site, the common biologically relevant signals arising during longer-term evolution can be captured in relatively low dimension.

### 1.5 Visualization of evolution along an empirical fitness landscape

In RNA virus evolution, adaptation to new environments can often be attributed to single or few new mutations that become fixed in the population. Experimental evolution in the lab and convergent evolution in the field suggest that short term evolution may be of relatively low dimension, as supported by our findings. If so, then these initial movements in sequence space may be inherently predictable, provided a robust genotype-phenotype map could be generated. This connection between sequence space and fitness is most naturally illustrated as a fitness landscape, where fitness is shown as a function of location in sequence space. However, reconstructing such landscapes from empirical data has been challenging. To evaluate the ability of DISSEQT to correctly generate and visualize fitness landscapes, we first empirically measured the relative fitness of the wild type and SynSyn virus populations described above in a direct competition assay against a neutral, genetically marked competitor^26;29^ (data available in Synapse). The visualization (Figure 5, top panel) builds upon a 2d representation of sequence space, but using only the first two components from the SMSSVD representation is not sufficient since it ignores all other relevant signals in the data. Nonlinear dimension reduction by Isomap was used to distort sequence space such that the notion of closeness is respected, taking all signals into account. Fitness was then added as the third dimension, interpolated by the Gaussian Kernel Smoother predictor (performance measured in Figure 6, top panel). The figure shows the dynamics of viral clouds corresponding to each viral lineage evolving over time. The wild type lineage (black) occupied the centermost area of the landscape, surrounded by the other lineages. In general, wild type populations occupied high fitness regions of the landscape, with some variability. This observation confirmed that wild type virus is well adapted to the growth conditions used in these experiments, and should tolerate perturbations in the system, such as increases in mutational load. The green SynSyn lineage displayed the most dramatic fitness differences, reaching both very high and very low areas, whereas the blue SynSyn lineage showed a stable, plateau-like behavior without any significant drops in fitness. Finally, the red SynSyn lineage was stuck in an area of the fitness landscape with-out any fitness peaks. Indeed, the red lineage was shown to be attenuated in vivo, and unable to reach pathogenic outcomes available to wild type virus; while the blue lineage was shown to be more mutationally robust^26^. Supplementary Figure 4 shows the same fitness landscape, but with samples colored by mutagen. Importantly, the data show that 2d reconstruction of sequence space by nonlinear dimension reduction can adequately reconstruct a fitness landscape that captures the expected biological behavior of similar, yet different viral lineages.

**Figure 5:**
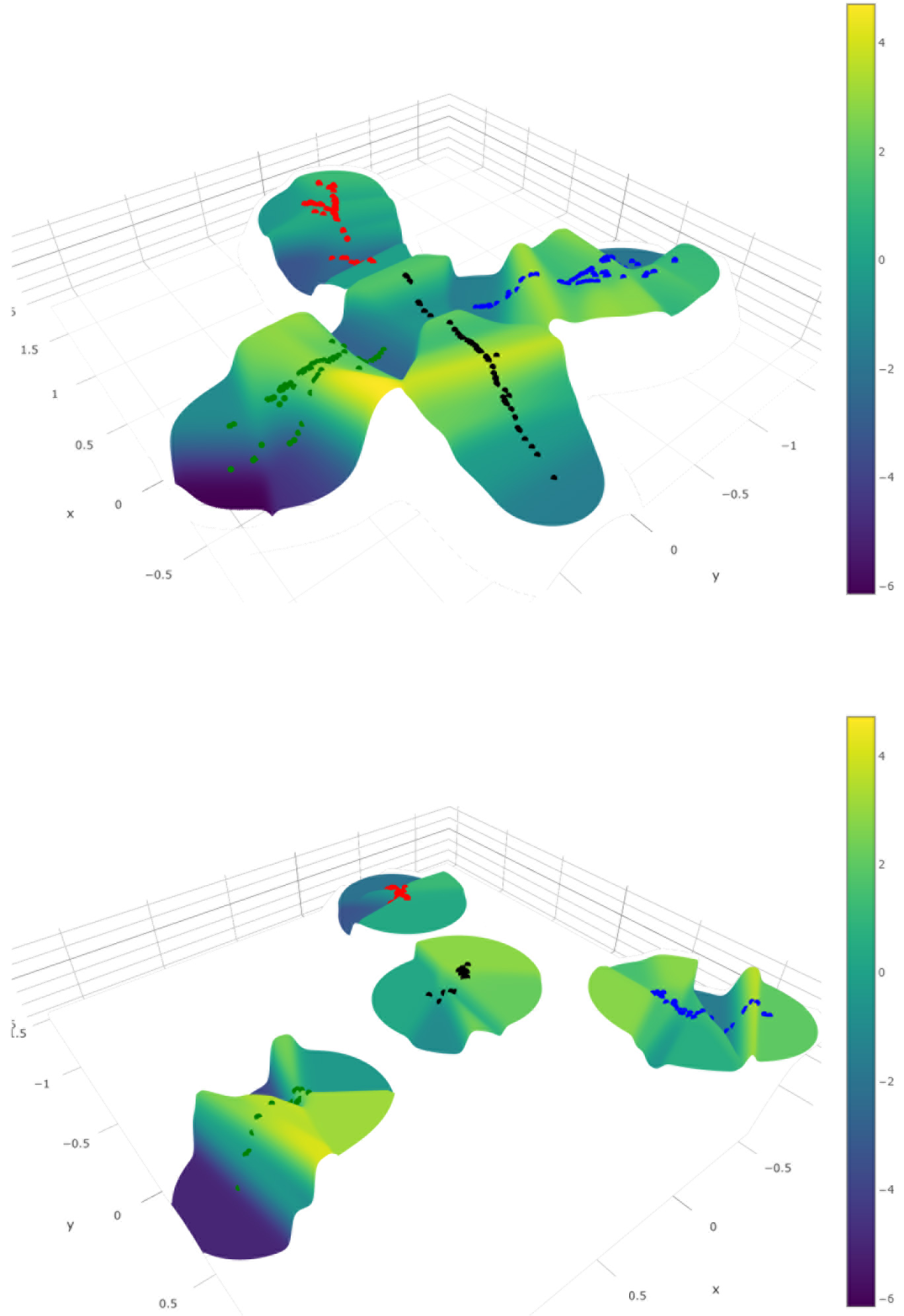
Top: Fitness landscape visualization of the SynSyn data set. Bottom: The same fitness landscape, constructed from consensus data only. Samples are colored by lineage 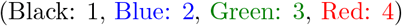.

**Figure 6:**
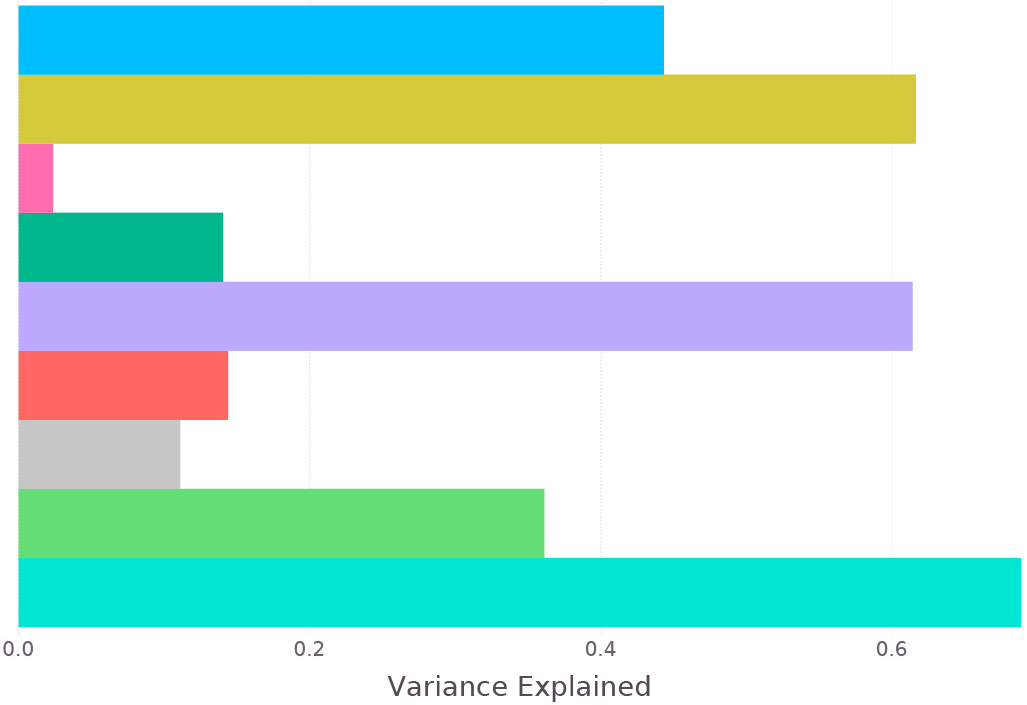
Comparison between different fitness predictors. Gaussian Kernel Smoother Predictors: Isomap 2d 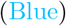, SMSSVD 13d 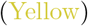, Consensus Isomap 2d 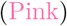 and Consensus 13d 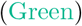. Nearest Neighbor Predictors: SMSSVD 13d 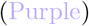 and Consensus 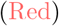. Group Predictors: Lineage/Mutagen 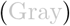, Lineage/Dose 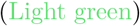, Lineage/Mutagen/Dose 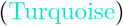.

### 1.6 Prediction of phenotype from genotype requires the input of minority variants

A prime goal in developing faithful representations of sequence space is the potential to assign phenotypes to known genotypes, and ultimately predict the phenotypes of new genotypes. For rapidly evolving populations, the presence of minority variants has been shown to contribute to phenotype, but this is not normally taken into account in genotype-phenotype mapping. Indeed, when the fitness landscape described above was reconstructed using only consensus sequence data, the landscape is considerably collapsed (Figure 5, bottom panel).

We thus evaluated the relevance of our sequence space reconstructions (after noise reduction) in their ability predict virus fitness, a quantitative parameter often used to describe phenotype. The performance of different fitness models was compared (Figure 6). Predicting fitness is inherently difficult. Thus, to get a baseline for the optimal performance that could be achieved, we used group-based predictors that rely on sample conditions, rather than deep sequencing data. The fine-grained group predictor using Lineage, Mutagen and Dosage accurately described the sample conditions (Figure 6, turquoise bar). In other words, when these three groupings are known for a sample, the prediction is over 69% accurate. When only lineage and dose were considered, prediction was 36% accurate, and if only lineage and mutagen were known, accuracy dropped to 11%. For the landscape predictors based on the 2d Isomap, accuracy was 44%. SMSSVD, on the other hand, which uses 13d reaches predictability of 62% and 61% from landscape or nearest neighbor predictors. The data revealed that while 2d Isomap performs well for visualization, prediction is best achieved when more components are incorporated. Importantly, when either Isomap or PCA is performed solely on consensus sequences, prediction fails (2% and 14%, respectively). Furthermore, the performance of the SMSSVD predictors compared well to the predictor based on experimental conditions (Lineage, Mutagen and Dose), the closest we have to a gold standard. In summary, the predictors based on our proposed sequence space representation vastly outperformed the consensus-based predictor. The data thus confirm that consensus sequencing of a viral population is not enough to understand its properties and cannot accurately predict its phenotype.

## 2 Discussion

High-throughput sequencing is replacing more classic sequencing methods in microbiology, especially in studying RNA viruses, where every nucleotide can be easily covered with extreme depth. This has increased and renewed interest in better characterizing RNA virus populations to take into account their variability, particularly when trying to identify differences between clinical or experimental samples that have no significant differences in consensus sequence, yet present different phenotypes. Recent works show that indeed, most sites along a genome generate mutants at very low frequency. Following passage of poliovirus in cell culture, Acevedo et al.^5^ identified an average of 16,500 variants, the equivalent of ~ 74% of all possible variant alleles in each passaged sample. Similarly, the previous analysis of the Coxsackie virus B3 wildtype populations described in more detail here, identified variant alleles in 65-80% of the sequenced regions^4^.

Despite the increasing accessibility of sequencing technology, we still lack the computational tools to use this data to its full potential. For instance, while an exhaustive list of variants can be generated per sample, to differentiate between similar, yet different, populations most studies have had to settle with using very basic mean measures such as Shannon entropy or mean variance. At best, these were followed up by a more targeted (and biased) focus on the few alleles suspected or known to be involved in the biological question being addressed.

A pre-existing obstacle to developing these tools was the uncertainty as to the size and dimensionality of sequence space actually occupied by evolving microbial populations. Mathematical sequence space is vast, even for the small genomes of RNA viruses. Theoretically, the high mutation rates of RNA viruses could reach a large amount of this space, questioning whether the evolution of these microbes could be inherently predictable. However, it is clear that biological constraints prohibit this from occurring, as most mutations will affect form or function and will not accumulate under strong purifying selection. In vivo and in vitro experimental evolution studies performed in independent replicates reveal that under a constant environment, the same set of mutations tend to emerge. This suggests that the sequence space available to a virus is indeed more limited, determined by its current genome sequence, raising the possibility that evolutionary trajectories may therefore be predicted at least in the very short term (the next one or few mutational steps). Fitness landscapes help understand what neighboring populations might represent distributions of genotypes of equal or increasing fitness and which regions define populations of lower fitness. Knowledge of fitness in the vicinity of a current population may help determine the most likely paths that will be taken during the evolution of the population. While this goal may seem lofty for large genomes, the small and highly constrained genomes of RNA viruses may be more amenable to such an exercise.

We have shown how the DISSEQT pipeline, using distribution-based modeling of complex, evolving microbial populations, can uncover many different genotypic and phenotypic patterns without needing a priori hypotheses of which genetic alleles are to be studied. Importantly, the robust dimension reduction methods performed here have successfully separated biologically relevant signals from sequencing-related error and other noise, identifying key characteristics of the quasispecies cloud that drive evolution. This accentuates that global properties, like the shape of the quasi-species cloud, are significant when trying to predict viral evolution. Sequencing error has long been an issue with characterizing microbial diversity and identifying true SNVs. Despite the presence of sequencing error, DISSEQT succeeded in finding structure in sequence space, made clear by the co-localization of populations subjected to similar environmental conditions and by accurate fitness predictions and fitness landscapes constructed on top of the sequence space representation.

Applied here, DISSEQT analysis has provided two key pieces of information regarding evolving RNA virus populations. First, that the biologically ‘relevant’ sequence space occupied by such populations is of intrinsically low dimension. In both data sets presented here, the SynSyn viruses that were manipulated to present discrete biological signals and the High-, Low- and Wildtype fidelity viruses evolving naturally to generate discrete differences in variant composition, the genetic signatures of biological interest were segregated and identified within an intrinsic space of very low dimension (10-20). Second and most importantly, we show that reliable prediction of phenotype from genotype requires the input of minority variants, underscoring the importance of studying RNA viruses, and perhaps other microbial organisms, as a population rather than as a single reference sequence.

At the core of our model is the representation of a population as a measure over a suitable genetic space. Using traditional bulk experimental techniques, averaging is performed already in the sampling and measuring steps of the management and analysis protocol; often resulting in relatively robust and normally (or log-normally) distributed data, well adapted for well-established statistical and machine learning analysis and visualization techniques.

We have shown that by directly modeling and representing the distribution at each genetic loci of all measurable minority variants, followed by model reduction, we get low dimensional and robust models that capture the interaction between minority variants and, by coupling it with phenotypic measurements, make it possible to follow and predict trajectories in genotype-phenotype space. It opens up for extending the sequence space models presented in this work to situations where heterogeneity of populations can be hypothesized to be an important aspect that can be measured in a direct manner. In particular, data coming from single cell sequencing have more variability, more artifacts and often complex distributions^13^ and distribution-based modeling can be envisioned to be viable and provide a natural and biologically accurate representation of the data.

Cancer growth, fundamentally different in origin from viruses and bacteria, may still be usefully described in terms of similar evolutionary processes^30^. Larger genomes and frequent structural variation, such as chromosomal aberrations^31^ and fused genes^32^ does, however, make the situation more complex and further work is needed to adapt the sequence space modeling for these circumstances. The challenges lie in incorporating structural variants into to the underlying space in a way that preserves biological similarity and is feasible to infer from the data. A possible starting point for cancer data is to restrict the analysis to a chosen set of interesting genes that do not exhibit any structural variation, thus simplifying the collection of deep sequencing data and providing an easy fit to the sequence space models we propose.

## 3 Methods

### 3.1 Reproducible and Traceable Analysis

Traceability in the DISSEQT pipeline is provided by integration with the collaborative science platform Synapse. Every result produced by DISSEQT can be traced back all the way to the original data files using the Synapse *provenance graph*, which describes the actions taken for every analysis step and connects input to output data. Sharing settings in Synapse makes it possible to open up the entire analysis to the public, but keeping sensitive data and unfinished analyses private if necessary. The analysis steps are self-contained in the sense that all data required to produce the output is downloaded from Synapse as needed. Hence, every analysis step can be reproduced locally by anyone executing the same actions. By changing parameters or making other changes, the impact of performing the analysis in a different manner can be investigated by others. Rerunning the entire analysis is also possible in this way. Furthermore, the analysis can be adapted to new data sets, such that the results can reproduced from new biological data.

### 3.2 Iterative Alignment

Alignment of sequenced reads to a reference genome was done with BWA-MEM ^33^. The choice of alignment tool is not critical, but the same one should be used for all samples to get a consistent analysis. After alignment, the consensus sequence of the aligned sample is computed. If the consensus differs from the reference genome, the alignment starts over, now using the consensus as the new reference genome. This process is repeated until the consensus has converged. Iterative alignment combats an inherent problem that occurs when aligning to a reference genome - there will be a bias since reads that match the reference genome are easier to align, while reads that differ might be mapped incorrectly or cut off such that the variant is not included in the alignment. For variants at the majority level, iterative alignment thus ensures that more reads are mapped correctly, allowing for a better frequency estimate. Even more important is that the ability to detect minority variants in the vicinity of majority level variants is greatly improved, as the number of differences between reads containing the minority variant and the consensus will tend to be lower.

### 3.3 Quality Control

Generating deep sequencing data is a complex procedure with many steps performed, both for the experiment itself and to prepare the data for sequencing. The DISSEQT pipeline provides several ways to evaluate the data to make sure that it is of high quality. Before alignment, adapters and poor quality bases are trimmed from the ends of reads using fastq-mcf^34^. At the end of the iterative alignment procedure, consensus sequences are automatically generated for all samples. It is expected that the consensus sequence will be more similar to the reference used for the initial alignment iteration, than to any other reference used in the same sequencing run. If this is not the case, the sample is flagged as being mislabeled. Indels are also reported. Graphs showing the read coverage as a function of genome position are created. All samples in the same sequencing run (and using the same reference genome) are put in the same graph, making it possible to identify problems with low read coverage for certain samples or genomic regions at a glance. Samples with a low mean read coverage can be removed automatically from downstream analysis. What threshold to use depends on the experimental setup, but we recommend keeping only samples with a mean read coverage above 1000 for deep sequencing data. There are also tools in DISSEQT to remove samples that are suspected of being contaminated by other samples, identified by having a mixture of reads that are likely to originate from different reference genomes. The purpose of quality control is to validate that we are indeed studying what we set out to study. If a sample is showing unexpected patterns, in particular during quality control, we recommend that the aligned reads, the consensus sequence and any other measurements are inspected manually ensure that conclusions are not drawn from faulty data.

### 3.4 Haplotypes

Recovering the haplotype mix from a collection of short reads is a difficult, often ill-conditioned, and computationally intensive problem, but several software tools^35;36^ are available, also see^37;38^ for overviews. The dominant haplotypes and their frequencies do not, however, completely characterize the viral population, another important aspect is how dispersed the individual viruses are around these central haplotypes. V-Phaser^39^, V-Phaser 2^40^ and ShoRAH^35^ find phased variants, pushing down the detection limit by assuming that real variants (at nearby loci) tend to co-vary, while errors do not. Unfortunately, V-Phaser and ShoRAH does not scale well for large data sets and V-Phaser 2 requires paired-end reads. For the reasons above, we chose the simpler and more robust path of making maximum likelihood (ML) estimates of the variant frequencies at each position, based on base quality data.

### 3.5 Sequence Space Representation

The genomic composition of microbial populations can be represented by a positive measure over a suitable space. Let ∑ be an alphabet set, e.g. the set of nucleotides 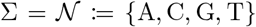, the set of codons 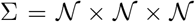 or the set of amino acids 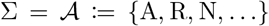. For the rest of this article, the set of codons will be used as the alphabet set, since the codons are closely connected to biological function and this choice doesn’t impose any assumptions about the relative importance of synonymous versus nonsynonymous changes. The set of codons is the natural choice for coding regions, to analyze non-coding regions, the set of nucleotides could be used instead. Now define *sequence space* ∑^*n*^ as the set of ordered sequences of length *n* over the alphabet ∑. Assuming that individual genomes in the population only differ by a finite number of point mutations (i.e. substitutions), the composition of the population is characterized by a positive measure over sequence space. The space of positive measures over sequence space will be denoted by 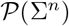.

Inference of the population composition can be intractable from sequencing data due to short reads and/or high error rates. Let 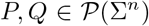 and define an equivalence relation such that *P* ~ *Q* iff

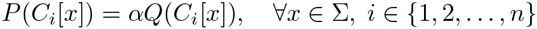

for some constant *α* ∈ ℝ^+^, where

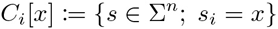

are the basic cylinder sets of ∑^*n*^. Hence, *P* relates to *Q* if they have the same allele frequencies at all positions. Inference for the equivalence class [*P*] from sequence data is possible even when *P* cannot be inferred since allele frequencies at different position can be estimated separately. The drawback is that minority variant linkage is lost. Each equivalence class [*P*] is naturally represented by the frequency matrix *p* ∈ ℝ^*n*×|∑|^ with *p*_*i*,*x*_ = *P*(*C*_*i*_[*x*])/*P*(∑^*n*^). Finally, the frequencies are transformed by *p* → log_2_(*p* + *α*), where α denotes the limit of detection, to give minority variants higher impact in the model. The log transformation emphasizes relative differences in frequencies between variants instead of absolute differences in frequencies between variants.

### 3.6 Sequence Space Inference

Maximum likelihood estimation was used to infer the codon frequencies at any given position, using all reads overlapping that position, based on a multinomial model with noisy observations. At a given locus, let ***θ*** = (*θ*_1_, *θ*_2_,…, *θ*_64_) be the frequencies in the population for the 64 different codons, with *θ*_*i*_ ≥ 0 for all *i* and ∑_*i*_*θ*_i_ = 1. Consider a read and the fragment the read is sequenced from. Now let *x* be the observed codon in the read and *z* the unknown codon in the original fragment, then

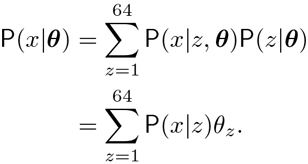

We model P(*x*|*z*) using the quality scores of the bases in the codon. If *ϵ*_1_, *ϵ*_2_, *ϵ*_3_ are the probabilities of a read error at bases 1, 2 and 3 in the codon and *y*^*k*^ is the base at position *k* in a codon *y*, then

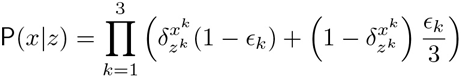

where 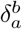 is the Kroenecker delta, the errors are thus assumed to be independent between bases in the codon and read errors are assumed to be equally likely to result in any of the other 3 bases. Assuming independent reads, the probability of the observations is

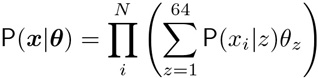

with observed codons ***x*** = (*x*_1_, *x*_2_,…, *x*_*N*_) from reads 1 to *N*. The log-likelihood is thus

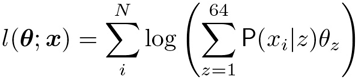

which is maximized numerically.

We noted that in our high read coverage data, bases with low-quality Phred scores tended to be biased toward certain nucleotide errors. Thus, we chose to not trust bases with a Phred score below 30. This was done by setting the ɛ_k_ of such nucleotides to 0.75, giving them no influence. Reads were excluded from the analysis if they caused the ML optimization problem to be underdetermined (e.g. when observing two reads with codons AAA and xAT respectively, where x means that the nucleotide is unknown, xAT is dropped since only the sum of the frequencies for AAT, CAT, GAT and TAT can be determined).

### 3.7 Limit of Detection

True minority variants can be hard to separate from sequencing errors. And in both cases, we expect the frequencies to be different depending on the nucleotide neighborhood and other factors^41^. A key difference is however that there are two sets of observations of the sequencing errors since the reads originating from the forward and reverse strands have different nucleotide neighborhoods for any given codon site. Indeed, for each sample, the codon frequencies from the two strands are expected to be approximately equal for true minority variants, something which is much less likely for sequencing errors. The differences in sequencing error behavior depending on the context thus leads us to estimate the limit of detection α separately for each locus and codon. For a given locus, the samples are grouped by run and consensus codon, to get similar sequencing errors across the samples in each group. Fix a codon and let ***f*** and ***r*** be two vectors where *f*_i_ and *r*_i_ are the inferred codon frequencies using reads from only the forward and reverse strands respectively, for sample *i* in the group. To limit the impact of sequencing errors on the downstream analysis, the transformed frequencies should be approximately equal, i.e. give a low value of the norm

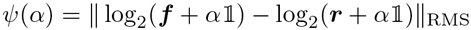

where log_2_ acts elementwise and 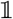 is a vector of all ones. Now define the limit of detection

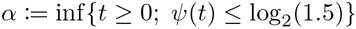

The infimum exists since *ψ* is continuous and *ψ*(*t*) → 0 as *t* → ∞. The threshold log_2_(1.5) is chosen such that if we have a single sample with *f*_1_ = *x* and *r*_1_ = 0, then *α* = 2*x*. Furthermore, *ψ* is a strictly decreasing function and *α* can thus be found by the bisection method or other root-finding methods. Finally we choose a conservative estimate of the limit of detection *α*_*c,x*_, for codon *c* at locus *x*, by taking the highest limit of detection estimated from the different sample groups 1, 2,…,*G*,

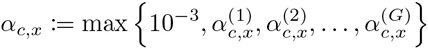

with upper indices denoting the sample group and where 10^−3^ is a commonly accepted lower limit of detection for sequencing data ^42^.

### 3.8 Dimension Estimation using Talus Plots

The Talus Plot provides a visualization of how the contents of a data set spread out across different dimensions and is designed to make it as easy as possible to make a qualitative estimate of the number of dimensions needed to capture all signals that stand out above the background noise. In Supplementary Methods, we show how predictable aspects of the background noise can be used to discern signals from noise. In brief, when the Talus Plot has “settled”, with small variations around a low mean, then the noise can be expected to be dominant.

### 3.9 SMSSVD

SubMatrix Selection Singular Value Decomposition (SMSSVD)^25^ is a parameter-free dimension reduction technique designed for the reconstruction of multiple overlaid low-rank signals from a data matrix, corrupted by noise. It is ideal for exploratory analysis of complex data, where different signals are spread out over different (possibly overlapping) subsets of variables, by limiting the influence of noise in variables that are not contributing to the signal. One of the major benefits of SMSSVD is its ability to detect signals with a low signal-to-noise ratio. SMSSVD shares many relevant properties with SVD, in particular orthogonality between components and the ability to extract variable loadings. The DISSEQT pipeline uses SMSSVD for noise reduction of the sequence space representation, since the number of variables is very large and we are trying to recover all signals that can be detected, not only the strongest one. Before applying SMSSVD, the data matrix is centered.

### 3.10 Fitness Landscapes

Fitness landscapes, an important kind of genotype-phenotype map, are used to illustrate the connection between sequence composition and fitness of organisms. Here we show how a fitness landscape can be generated entirely from empirical data. Given a d-dimensional representation of sequence space, i.e. a set of sample points *x*^(*i*)^ ∈ ℝ^*d*^, *i* = 1,…, *N* with corresponding fitness values *y*^(*i*)^ ∈ ℝ, we want to reconstruct a surface *f* : ℝ^*d*^ → ℝ such that

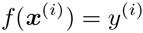

In practice however, we cannot expect a perfect fit of the surface. Differences in fitness between sample points that are close in the low dimensional representation will be difficult to capture. Furthermore, measurement noise will impact the reproducibility of the surface. To get a robust fitness landscape, we use a Gaussian Kernel Smoother^43^ and select the kernel width *σ* by cross-validation (repeated random subsampling). That is, we numerically find

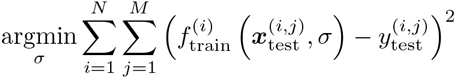

where the data is randomly divided into a train and a test data set for each iteration *i* and

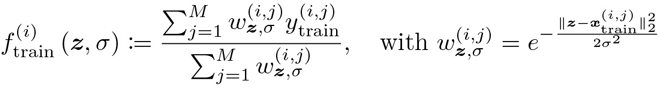

### 3.11 Fitness Evaluation

The Gaussian Kernel Smoother (fitness landscape) predictors are evaluated in comparison to other fitness predictors. Nearest Neighbor predictors uses the fitness of the closest sample in sequence space as the prediction and can be used for different sequence space models. In case of ties, the prediction is taken as the average over the tied samples. Group-based predictors use a predetermined grouping of the samples, predicting fitness as the average fitness among samples in the same group, and do not use sequence data at all.

Model accuracy for a predictor *f* is measured by fraction of variance explained,

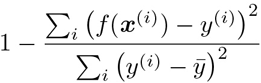

where *x*^(*i*)^ is the representation of sample *i* used by the predictor, *y*^(*i*)^ is the fitness of sample *i*, *ȳ* the mean fitness over all samples and the second term in the expression is the variance of the residuals divided by the total variance. The models are evaluated by leave-one-out cross validation. The kernel widths for the Gaussian Kernel Smoother predictors are estimated separately for each problem instance to avoid influence from the left-out sample.

### 3.12 Variable Selection

We use SPC (Sparse Principal Components) ^44^ for variable selection, after noise reduction by SMSSVD. SPC adds a variable-side *L*_1_ (lasso) constraint to a formulation of SVD as an optimization problem, forcing sparsity by ensuring that many variables are 0 at the optima. The optimization problem is then solved for one component at a time, using an iterative algorithm. However, since the optimization problem is not necessarily convex, the algorithm might converge to a local optima. To reduce the impact of this problem, and to ensure that the singular values are declining, we suggest an extension of the algorithm. It can be shown that if a component has a larger singular value than a previous one, then this solution is guaranteed to be a better starting guess for the optimization problem for the previous component. By rolling back and restarting the optimization at the previous component, we get closer to the globally optimal solution and make sure that the singular values are declining.

### 3.13 Nonlinear Dimension Reduction

By dimension reduction, we aim to identify the parts of sequence space that are explored by the samples. Linear dimension reduction techniques, like SMSSVD, are useful because they make very few assumptions about the structure of the data. Although there is no reason to believe that the underlying manifold is linear, the complexity that is necessary for biological systems is indeed often caused by nonlinearities, linear methods can still capture nonlinear patterns if the dimension is sufficiently high ^45^. However, to get an informative visualization in just two or three dimensions, nonlinear dimension reduction is needed for complex data sets.

We apply Isomap^46^ to the data set after the noise reduction by SMSSVD (and the optional variable selection). Nonmetric multidimensional scaling us-ing Kruskal’s stress criterion^47^ was used rather than classical multidimensional scaling in the final step of the Isomap algorithm. This distorts the underlying space by expanding local structure that would otherwise be too small to notice, giving some importance to weaker signals in the data.

### 3.14 Time Series

The evolution of a population over time is described by a curve ***p***(*t*) in sequence space. In practice, we can only measure the values of a curve ***p***(*t*) at discrete time points, and the measurements are subjected to noise. As the first step of noise reduction, a 3-point median filter over time is applied to the sequence space representation, to robustly reduce the impact of noise spikes. Following the noise-reduction, the curve ***p***(*t*) is reconstructed in the *d*-dimensional representation of sequence space, as a piecewise linear curve connecting the data points. Then, each curve is reparameterized by arc length *s*, starting at *s* = 0 for *t* = 0, since differences in mutation rates can cause the population to move at different speeds through sequence space.

The sequence space representation in terms of variables (variants) is time-invariant, but it is nevertheless important to see how different parts of sequence space are explored as the replicates move. Let *σ* be the first singular value, with corresponding left and right singular vectors ***u*** and ***υ***, after dimension reduction of a matrix *X* by SMSSVD, SVD or SPC, then *σ* can be decomposed as a sum over variables and samples,

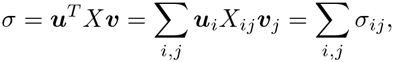

where *σ*_*ij*_ := ***u***_*i*_*X*_*ij*_***v***_*j*_ quantifies the importance of variable *i* and sample *j* for this component. By linear interpolation, this can be extended to 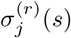, for intermediate values of the curve parameter *s* for replicate *r*. The contribution of variable *j* at *s* is measured by 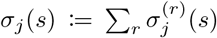 and *σ* := ∑_*j*_ *σ*_*j*_(*s*) describes the importance of the first principal component at *s*. Plotting *σ*(*s*) and *σ*_*j*_(*s*) along with the replicates, thus aid understanding of the dynamics. The definitions naturally extend to multiple components.

### 3.15 Bifurcations

We define the time of bifurcation *β*(***p***, ***q***), between two curves ***p***(*t*) and ***q***(*t*) as the similarity measure

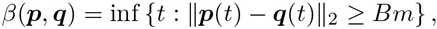

that is, the first point in time at which the distance between ***p***(*t*) and ***q*** is above a threshold. Here, *B* is a chosen threshold and *m* a normalization constant chosen to make the expression scale-invariant. If ***p***^(*i*)^(*t*) is defined for *t* ∈ [0, *T*_*i*_] and *T*_*ij*_ := min(*T*_i_, *T*_*j*_), then the mean distance over time between curves *i* and *j* is

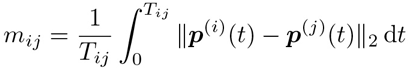

and we let 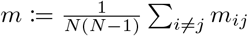, the mean over all pairs of the *N* curves. Average linking hierarchical clustering, based on the time of bifurcation similarity scores, naturally extends the concept to clusters of samples, giving recursive cluster splits and a cluster similarity score equal to the time of bifurcation at each split. For piecewise linear curves, *m* and *β*(***p***^(*i*)^, ***p***^(*j*)^) can be computed analytically.

## Acknowledgements

This work is sponsored by the Defense Advanced Research Projects Agency INTERCEPT program managed by Dr. Jim Gimlett, and administered though DARPA contract HR00111720023: the content of the information does not necessarily reflect the position or the policy of the Government and no official endorsement should be inferred.

## References

[1] Walter Fiers, Roland Contreras, Fred Duerinck, Guy Haegeman, Dirk Is-erentant, Jozef Merregaert, W Min Jou, Francis Molemans, Alex Raeymaekers, A Van den Berghe, et al. Complete nucleotide sequence of bacteriophage ms2 rna: primary and secondary structure of the replicase gene. Nature, 260(5551):500–507, 1976.

[2] Robert D Fleischmann, Mark D Adams, Owen White, Rebecca A Clayton, Ewen F Kirkness, Anthony R Kerlavage, Carol J Bult, Jean-Francois Tomb, Brian A Dougherty, Joseph M Merrick, et al. Whole-genome random sequencing and assembly of haemophilus influenzae rd. science, pages 496–512, 1995.

[3] Claire M Fraser, Jeannine D Gocayne, Owen White, Mark D Adams, Rebecca A Clayton, Robert D Fleischmann, Carol J Bult, Anthony R Kerlavage, Granger Sutton, Jenny M Kelley, et al. The minimal gene complement of mycoplasma genitalium. science, pages 397–403, 1995.

[4] Antonio V Bordería, Ofer Isakov, Gonzalo Moratorio, Rasmus Henningsson, Sonia Agüera-González, Lindsey Organtini, Nina F Gnädig, Hervé Blanc, Andrés Alcover, Susan Hafenstein, et al. Group Selection and Contribution of Minority Variants during Virus Adaptation Determines Virus Fitness and Phenotype. PLOS Pathogens, 2015.

[5] Ashley Acevedo, Leonid Brodsky, and Raul Andino. Mutational and fitness landscapes of an rna virus revealed through population sequencing. Nature, 505(7485):686, 2014.

[6] Marco Vignuzzi, Jeffrey K Stone, Jamie J Arnold, Craig E Cameron, and Raul Andino. Quasispecies diversity determines pathogenesis through cooperative interactions in a viral population. Nature, 439(7074):344–348, 2006.

[7] Katherine S Xue, Kathryn A Hooper, Anja R Ollodart, Adam S Dingens, and Jesse D Bloom. Cooperation between distinct viral variants promotes growth of h3n2 influenza in cell culture. Elife, 5:e13974, 2016.

[8] J Maynard Smith and George R Price. The logic of animal conflict. Nature, 246(5427):15–18, 1973.

[9] Johannes G Reiter, Ayush Kanodia, Raghav Gupta, Martin A Nowak, and Krishnendu Chatterjee. Biological auctions with multiple rewards. In Proc. R. Soc. B, volume 282, page 20151041. The Royal Society, 2015.

[10] Charles Gawad, Winston Koh, and Stephen R Quake. Single-cell genome sequencing: current state of the science. Nature reviews. Genetics, 17(3): 175, 2016.

[11] Jeffrey M Perkel. Single-cell sequencing made simple. Nature, 547(7661): 125, 2017.

[12] Valentine Svensson, Kedar N Natarajan, Lam-Ha Ly, Ricardo J Miragaia, Charlotte Labalette, Iain C Macaulay, Ana Cvejic, and Sarah A Teichmann. Power analysis of single cell rna-sequencing experiments. bioRxiv, page 073692, 2016.

[13] Rhonda Bacher and Christina Kendziorski. Design and computational analysis of single-cell rna-sequencing experiments. Genome biology, 17(1):63, 2016.

[14] Alistair B Russell, Cole Trapnell, and Jesse D Bloom. Extreme heterogeneity of influenza virus infection in single cells. bioRxiv, page 193995, 2017.

[15] Desmond G Higgins. Sequence ordinations: a multivariate analysis approach to analysing large sequence data sets. Computer applications in the biosciences: CABIOS, 8(1):15–22, 1992.

[16] Jiajie Zhang, Amir M Mamlouk, Thomas Martinetz, Suhua Chang, Jing Wang, and Rolf Hilgenfeld. Phylomap: an algorithm for visualizing relationships of large sequence data sets and its application to the influenza a virus genome. BMC bioinformatics, 12(1):248, 2011.

[17] Sewall Wright. The roles of mutation, inbreeding, crossbreeding, and selection in evolution, volume 1. na, 1932.

[18] Stuart A Kauffman and Edward D Weinberger. The NK model of rugged fitness landscapes and its application to maturation of the immune response. Journal of Theoretical Biology, 141(2):211–245, 1989.

[19] Michael C Whitlock and Denis Bourguet. Factors affecting the genetic load in drosophila: synergistic epistasis and correlations among fitness components. Evolution, 54(5):1654–1660, 2000.

[20] Jennifer A Collins, M Gregory Thompson, Elijah Paintsil, Melisa Ricketts, Joanna Gedzior, and Louis Alexander. Competitive fitness of nevirapine-resistant human immunodeficiency virus type 1 mutants. Journal of virology, 78(2):603–611, 2004.

[21] Roger D Kouyos, Gabriel E Leventhal, Trevor Hinkley, Mojgan Haddad, Jeannette M Whitcomb, Christos J Petropoulos, and Sebastian Bonhoeffer. Exploring the complexity of the hiv-1 fitness landscape. 2012.

[22] Christof K Biebricher and Manfred Eigen. What is a quasispecies? In Quasispecies: Concept and Implications for Virology, pages 1–31. Springer, 2006.

[23] David Seifert, Francesca Di Giallonardo, Karin J Metzner, Huldrych F Günthard, and Niko Beerenwinkel. A framework for inferring fitness landscapes of patient-derived viruses using quasispecies theory. Genetics, 199 (1):191–203, 2015.

[24] Kenneth A Stapleford, Lark L Coffey, Sreyrath Lay, Antonio V Bordería, Veasna Duong, Ofer Isakov, Kathryn Rozen-Gagnon, Camilo Arias-Goeta, Herve Blanc, Stéphanie Beaucourt, et al. Emergence and transmission of arbovirus evolutionary intermediates with epidemic potential. Cell host & microbe, 15(6):706–716, 2014.

[25] Rasmus Henningsson and Magnus Fontes. SMSSVD – SubMatrix Selection Singular Value Decomposition. ArXiv e-prints, October 2017.

[26] Gonzalo Moratorio, Rasmus Henningsson, Cyril Barbezange, Lucia Carrau, Antonio V Bordería, Hervé Blanc, Stephanie Beaucourt, Enzo Z Poirier, Thomas Vallet, Jeremy Boussier, et al. Attenuation of RNA viruses by redirecting their evolution in sequence space. Nature microbiology, 2:17088, 2017.

[27] Stéphanie Beaucourt, Antonio V Bordería, Lark L Coffey, Nina F Gnädig, Marta Sanz-Ramos, Yasnee Beeharry, and Marco Vignuzzi. Isolation of fidelity variants of rna viruses and characterization of virus mutation frequency. Journal of visualized experiments: JoVE, (52), 2011.

[28] Magnus Fontes and Charlotte Soneson. The projection score – an evaluation criterion for variable subset selection in PCA visualization. BMC bioinformatics, 12(1):307, 2011.

[29] P Carrasco, JA Darós, P Agudelo-Romero, and SF Elena. A real-time rt-pcr assay for quantifying the fitness of tobacco etch virus in competition experiments.Journal of virological methods, 139(2):181–188, 2007.

[30] Michael R Stratton, Peter J Campbell, and P Andrew Futreal. The cancer genome. Nature, 458(7239):719, 2009.

[31] Douglas Hanahan and Robert A Weinberg. Hallmarks of cancer: the next generation. cell, 144(5):646–674, 2011.

[32] Henrik Lilljebjörn, Rasmus Henningsson, Axel Hyrenius-Wittsten, Linda Olsson, Christina Orsmark-Pietras, Sofia Von Palffy,Maria Askmyr, Marianne Rissler, Martin Schrappe, Gunnar Cario, et al. Identification of etv6-runx1-like and dux4-rearranged subtypes in paediatric b-cell precursor acute lymphoblastic leukaemia. Nature communications, 7, 2016.

[33] Heng Li. Aligning sequence reads, clone sequences and assembly contigs with BWA-MEM. ArXiv e-prints, March 2013.

[34] Erik Aronesty. ea-utils: Command-line tools for processing biological sequencing data, 2011. URL https://github.com/ExpressionAnalysis/ea-utils.

[35] Osvaldo Zagordi, Arnab Bhattacharya, Nicholas Eriksson, and Niko Beerenwinkel. Shorah: estimating the genetic diversity of a mixed sample from next-generation sequencing data. BMC bioinformatics, 12(1):119, 2011.

[36] Mattia CF Prosperi and Marco Salemi. Qure: software for viral quasispecies reconstruction from next-generation sequencing data. Bioinformatics, 28 (1):132–133, 2012.

[37] Kerensa McElroy, Torsten Thomas, and Fabio Luciani. Deep sequencing of evolving pathogen populations: applications, errors, and bioinformatic solutions. Microbial informatics and experimentation, 4(1):1, 2014.

[38] Niko Beerenwinkel, Huldrych F Günthard, Volker Roth, and Karin J Metzner. Challenges and opportunities in estimating viral genetic diversity from next-generation sequencing data. Frontiers in microbiology, 3, 2012.

[39] Alexander R Macalalad, Michael C Zody, Patrick Charlebois, Niall J Lennon, Ruchi M Newman, Christine M Malboeuf, Elizabeth M Ryan, Christian L Boutwell, Karen A Power, Doug E Brackney, et al. Highly sensitive and specific detection of rare variants in mixed viral populations from massively parallel sequence data. PLoS Comput Biol, 8(3):e1002417, 2012.

[40] Xiao Yang, Patrick Charlebois, Alex Macalalad, Matthew R Henn, and Michael C Zody. V-phaser 2: variant inference for viral populations. BMC genomics, 14(1):674, 2013.

[41] Mark A DePristo, Eric Banks, Ryan Poplin, Kiran V Garimella, Jared R Maguire, Christopher Hartl, Anthony A Philippakis, Guillermo Del Angel, Manuel A Rivas, Matt Hanna, et al. A framework for variation discovery and genotyping using next-generation dna sequencing data. Nature genetics, 43(5):491–498, 2011.

[42] Sara Goodwin, John D McPherson, and W Richard McCombie. Coming of age: ten years of next-generation sequencing technologies. Nature Reviews Genetics, 17(6):333–351, 2016.

[43] Trevor Hastie, Robert Tibshirani, and Jerome Friedman. The elements of statistical learning, volume 2. Springer, 2009.

[44] Daniela M Witten, Robert Tibshirani, and Trevor Hastie. A penalized matrix decomposition, with applications to sparse principal components and canonical correlation analysis. Biostatistics, page kxp008, 2009.

[45] John Nash. The imbedding problem for riemannian manifolds. Annals of mathematics, pages 20–63, 1956.

[46] Joshua B Tenenbaum, Vin De Silva, and John C Langford. A global geometric framework for nonlinear dimensionality reduction. Science, 290 (5500):2319–2323, 2000.

[47] Joseph B Kruskal. Multidimensional scaling by optimizing goodness of fit to a nonmetric hypothesis. Psychometrika, 29(1):1–27, 1964.

